# Optimization of functional genetics tools for a model tetraploid Capsella bursa-pastoris, with focus on homoeolog-aware gene editing

**DOI:** 10.1101/2025.06.02.657020

**Authors:** D.O. Omelchenko, A.M. Barkovskaya, L.V. Omelchenko, A.V. Klepikova, A.A. Penin, M.D. Logacheva

**Affiliations:** Laboratory of Plant Genomics, Vavilov Institute of General Genetics, Russian Academy of Sciences, Moscow, Russia; Center for Molecular and Cellular Biology, Skolkovo Institute of Science and Technology, Moscow, Russia

**Keywords:** floral dip, CRISPR/Cas, *CO*, *SVP*, gRNA database

## Abstract

*Capsella bursa-pastoris* is a recent allotetraploid and a promising model for studying early consequences of polyploidy. One of the intriguing questions in polyploid research is how new functions arise from initially identical or nearly identical homoeologous genes. Functional genetics tools, including genetic editing, can help to understand this process, but they have not been developed for *C. bursa-pastoris* yet. We present here the results of our study aimed at filling this gap. In particular, we compared the efficiency of floral dip transformation in six accessions of *C. bursa-pastoris* representing distant populations. The Asian clade accession PGL0025 had the highest efficiency of transformation (∼1.1%). Comparison of *Agrobacterium tumefaciens* strains EHA105 and GV3101 (pMP90) showed that the latter is more effective. Also, we created a genome-wide gRNA database for all pairs of homoeologs of the PGL0001 accession of *C. bursa-pastoris* and integrated it into publicly available genome browser: https://t2e.online/igv_capsella_bursa-pastoris/. We assessed the possibility of differential editing for two pairs of homoeologous genes with high sequence similarity (>90%) both *in vitro* and *in silico*. Despite the test results that indicated off-target activity, we have succeeded in obtaining lines of plants with homozygous frameshift mutations in each of the homoeologs separately *in vivo*. We expect that these findings and resources will promote the use of *C. bursa-pastoris* as a model in functional genetics experiments, in particular, the studies of the fate of duplicated gene after polyploidization event.

## 2 Introduction

*Capsella bursa-pastoris*, commonly known as Shepherd’s purse, is an annual weed that belongs to the Brassicaceae family. It is a recent allotetraploid of hybrid origin, and unlike its parental species, it is ubiquitously distributed and has high morphological, physiological, and genetic diversity (Douglas et al. 2015; Wesse et al. 2021; Penin et al. 2024). This evolutionary success has drawn the attention of researchers from many areas of plant science, from ecologists to geneticists. Presumably, greater plasticity is enabled by diversification of functions between homoeologous genes (neo- and subfunctionalization) mediated by polyploidy. Due to its short life cycle and close relationship to the model species *Arabidopsis thaliana*, *C. bursa-pastoris* is a plausible model for studying the early consequences of polyploidy and its role in environmental adaptation and the emergence of morphological and physiological diversity (Nutt et al. 2006; Hintz et al. 2006; Slotte et al. 2007; Kryvokhyzha et al. 2016; Kasianov et al. 2017; Cornille et al. 2022).

To gain further insights into the genetic basis of these features, the use of functional genetic tools is extremely helpful. Among these tools, gene editing is the most powerful and promising (for a review, see (Gudmunds et al. 2022)). This approach potentially allows for the selective inactivation of homoeologous genes and exploration of resulting knockout plant phenotypes, highlighting the function of each homoeolog and possible interactions between them. Genetic editing can be used not only to create knockouts but also to perform allele replacement through the HDR pathway, introduce structural variations (e.g., (Beying et al. 2020)), and change methylation patterns of regulatory significance (Ghoshal and Gardiner 2021). Successful editing requires several steps, including the prediction of sgRNA sensitivity and specificity to avoid off-target effects (reviewed in (Cardi et al. 2023)). Polyploids present an additional challenge due to the presence of highly similar homoeologous genes, which are copies of genes originating from parental species after a hybrid speciation event. Most studies on genetic editing in polyploids aim to simultaneously target all homoeologs (e.g., (Wang et al. 2014; Aznar-Moreno and Durrett 2017; Singh et al. 2018; Zaman et al. 2019; Corkins et al. 2022)). This is reasonable when homoeologs act in an additive manner and complete inactivation of all copies is needed to achieve a desirable phenotype. This is, however, not always the case; for many polyploid plants, including *C. bursa-pastoris*, divergent expression patterns were found for many genes (Chaudhary et al. 2009; Pfeifer et al. 2014; Kasianov et al. 2017), implying more or less drastic divergence in function (sub- or neofunctionalization). This calls for homoeolog-aware editing, i.e., differential editing of each of the homoeologs. This might not always be feasible because of the high similarity of homoeologs. Here, we investigate the possibility of differential editing of homoeologs in *C. bursa-pastoris* as a model for recent allopolyploids using a recently published genome sequence. Another aspect of our study is the improvement of *Agrobacterium*-mediated transformation efficiency. To date, only one study (Bartholmes et al. 2008) has reported successful transformation (with an efficiency ranging from 0.1 to 0.4%) of *C. bursa-pastoris*. We expect that such low efficiency is not the limit. *C. bursa-pastoris* is a highly polymorphic species consisting of isolated groups that differ in a number of morphological and physiological characteristics. These groups form three large clades with distinct geographical distributions: European (EU), Middle Eastern (ME), and Asian (ASI). In the study (Bartholmes et al. 2008) only European accessions were tested. To address this gap, we conducted floral dip transformation test on a set of *C. bursa-pastoris* accessions representing the European, Middle Eastern, and Asian clades to identify those with the highest transformation efficiency.

## 3 Methods

### 3.1 Plant material and growth conditions

To evaluate the effectiveness of agrobacterial transformation we have chosen six accessions from our collection of *C. bursa-pastoris* (L.) Medik. ecotypes (Table 1). For each accession a single plant was grown in a growth chamber isolated from other *Capsella* plants. A single seed was randomly selected from a natural selfing progeny and grown in the same conditions. PGL0001, PGL0002, PGL0020, PGL0025, and PGL0066 were subjected to four rounds of self-pollination as described above, PGL0016 - to two rounds.

**Table 1.** Source of the *C. bursa-pastoris* accessions.

| Accession ID | Longitude | Latitude | Source region | Comment |
| --- | --- | --- | --- | --- |
| PGL0001 | 55.701936 N | 37.522078 E | Moscow, Russia | alias 'msu-wt' (Klepikova et al. 2021) |
| PGL0016 | 41.709234 N | 44.776140 E | Tbilisi, Georgia |  |
| PGL0002 | 55.701936 N | 37.522078 E | Moscow, Russia | alias 'lel4' (Klepikova et al. 2021)<br>(apetalous mutant) |
| PGL0025 | 25.139328 N | 102.743474 E | Kunming, China | alias 'KBG' |
| PGL0066 | 68.969386 N | 33.075942 E | Murmansk, Russia |  |
| PGL0020 | 31.773685 N | 35.226401 E | Jerusalem, Israel |  |

Prior to sowing, the seeds were stratified for 1 week at 4°C in pots filled with moist vermiculite. After stratification, the pots were transferred to flood and drain hydroponic systems (General Hydroponics Europe, France) supplied with GHE Terra Aquatica TriPart fertilizer solution (Gro + Micro HW + Bloom) at the manufacturers’ recommended concentrations. The plants were cultivated in a growth room under long-day conditions (16/8-h day/night light cycle, 15000 lux) at 21–22°C and 55–60% humidity.

### 3.2 Development of the genome-wide gRNA database for *C. bursa-pastoris* PGL0001

The spacer database was created using the crisprVerse v1.0.0 R library package (Hoberecht et al. 2022). The *C. bursa-pastoris* PGL0001 genome (GCA_001974645.2 GenBank) in FASTA format was converted to the appropriate format for the program using the R BSgenome library v.1.66.1 (Pagès 2022). Additionally, the genome annotation was standardized to the GFF3 format using the AGAT v1.2.0 script set (Dainat et al. 2023).

We have identified spacers for SpCas9 with a canonical “NGG” protospacer adjacent motif (PAM) in gene sequences using the R libraries crisprDesign v1.0.0 and crisprBase v1.2.0 according to the recommended pipeline of the authors of crisprVerse. To locate potential off-target sites, we performed iterative mapping of spacers across the entire genome using crisprBowtie v1.2.0 library. The spacers were evaluated for target editing efficiency using the crisprScore v1.2.0 library and the DeepHF (Wang et al. 2019) and DeepSpCas9 (Kim et al. 2019) algorithms, which are deep learning algorithms based on models trained on real gRNA activity data. To estimate potential off-target activity, we used the currently standard cutting frequency determination (CFD) metric (Doench et al. 2016). This metric calculates the probability of gRNA binding to a nontarget sequence by considering the location, number, and type of nucleotide substitutions between the spacer and protospacer.

The database was filtered to exclude spacers that are less likely to be effective in target gene knockout on the basis of their characteristics according to general guidelines for optimal gRNA design (Doench et al. 2016; Schindele et al. 2020). The following filters were applied:

1) Spacer GC composition should be between 20–80%;
2) Spacers should be located within 5–65% of the coding sequence (CDS) of the gene;
3) Spacers should not contain polyT sequences (four or more T nucleotides), as they may cause premature termination of sgRNA transcription;
4) The spacer should not have sequence complementarity with the sgRNA hairpin backbone of SpCas9 or itself to avoid disrupting the proper secondary structure of the sgRNA;
5) Spacers should not be located at overlapping sites of coding regions of different genes to avoid multitargeting activity;
6) The spacer cut site should be located in the CDS region of all the coding isoforms of a target gene;
7) DeepSpCas9 and DeepHF on-target scores should be at least 0.2, and the aggregated CFD off-target score (for all targets with a maximum of 3 mismatches) should be at least 0.2. We chose a low threshold to give users of the gRNA database a wider range of gRNAs to choose from, because target activity predictions based on models developed for human and animal cells are not entirely reliable for plants (Naim et al. 2020).

As a result, we obtained a comprehensive database of gRNA spacers for knockout of the annotated genes of *C. bursa-pastoris*. Using this database, we selected gRNAs for the knockout of *SVP_O*, *SVP_R*, *CO_O* and *CO_R* homoeologs (_R and _O denote the genes from the paternal subgenome, which is derived from *C. rubella*/*grandiflora* and maternal subgenome which is derived from *C. orientalis*, correspondingly).

### 3.3 DNA extraction, amplification and *in vitro* cleavage assay test

Primers for specific amplification of fragments of each homoeologous gene containing the corresponding gRNA target sequences were selected using the Primer-BLAST web service (Ye et al. 2012) using *C. bursa-pastoris* genome sequence as reference (GCA_001974645.2 NCBI GenBank). The characteristics of the selected primers are presented in Additional file 2, Table S1.

Plant DNA was extracted from fresh leaves of *C. bursa-pastoris* PGL0001 plants using SKYSuper Plant Genomic DNA Kit (SkyGen, Russia), and the concentration was measured with a Qubit 1X dsDNA High Sensitivity Kit (Invitrogen, USA) on a Qubit 3 (Invitrogen, USA). PCR was performed using Q5 Hot Start High-Fidelity 2X Master Mix (New England Biolabs) on a T-100 Thermal Cycler (Bio-Rad, USA). The PCR mixtures were purified using Agencourt AMPure XP (Beckman Coulter, USA) magnetic beads at a bead-to-sample v/v ratio of 0.9:1. The purified amplicon DNA was used in the subsequent SpCas9:sgRNA cleavage assay or Sanger sequencing for editing detection.

The SpCas9:sgRNA cleavage assay was performed using EnGen sgRNA Synthesis Kit, *S. pyogenes* (New England Biolabs, USA) and the EnGen Spy Cas9 NLS (New England Biolabs, USA) according to the manufacturer’s protocol. The results of the cleavage assay were evaluated using the Agilent High Sensitivity DNA Kit on a 2100 Bioanalyzer instrument (Agilent, USA).

The Sanger sequencing reads were subjected to semiautomated processing, trimmed to leave 200 bp upstream and downstream of the cleavage site and subsequently deposited in the GenBank database (Accession numbers PP711553-PP711570).

### 3.4 Design of binary vectors for homoeologous genes’ editing and transformation efficiency assessment in *C. bursa-pastoris*

For efficiency assessment of Agrobacterium-mediated transformation in *C. bursa-pastoris* a binary vector containing the pFAST-R marker (Shimada et al. 2010) and BASTA resistance expression cassettes was created with GoldenGate assembly method using plasmids from the Addgene MoClo Toolkit (Weber et al. 2011; Werner et al. 2012) and MoClo Plant Parts Kit (Engler et al. 2014) (Figure 1a).

**Fig.1.**
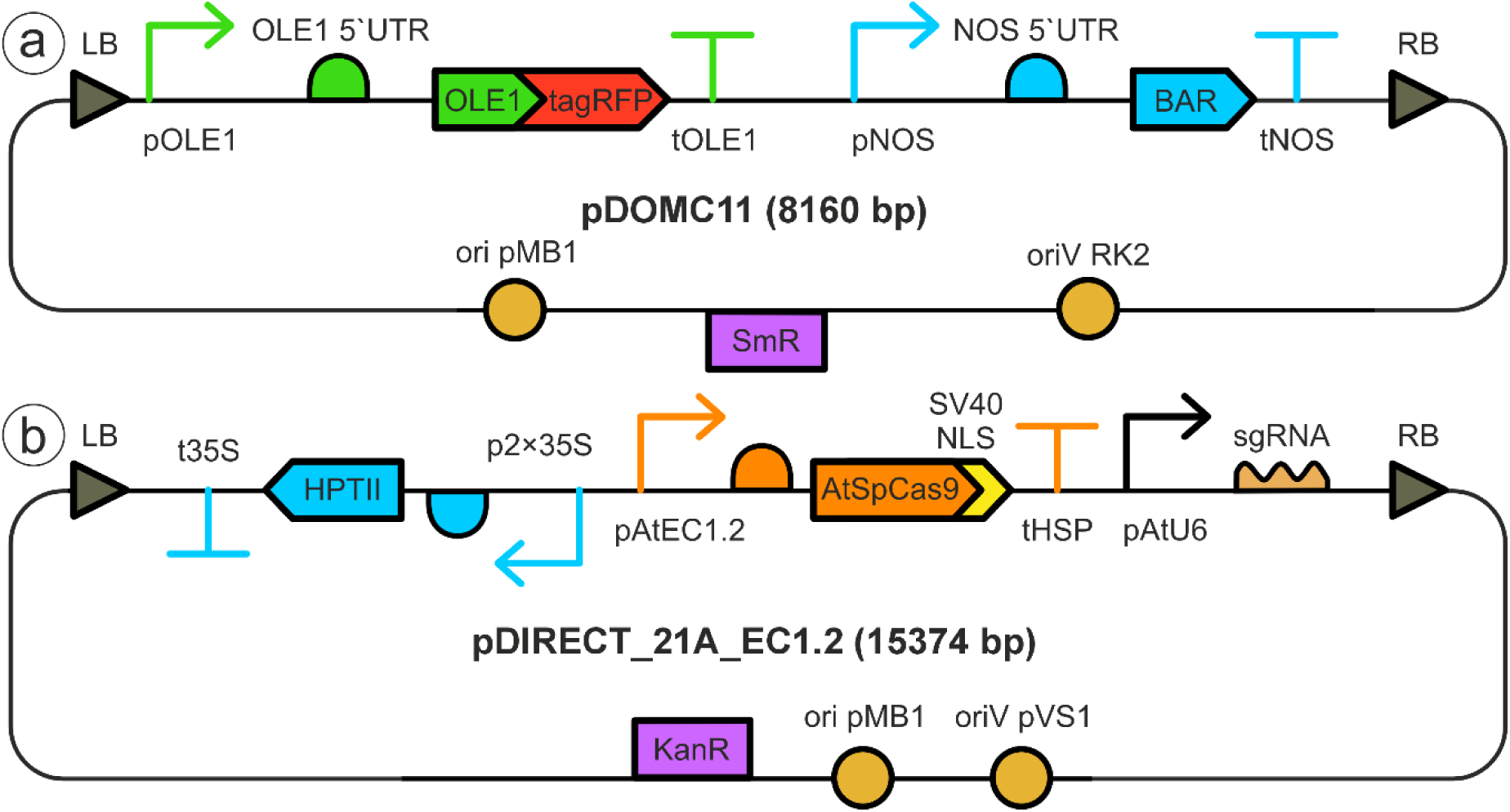
Schematic diagrams of the binary vectors used in the study. Vectors’ maps are presented in SBOL format (McLaughlin et al. 2020). (a) pDOMC11 binary vector, (b) modified pDIRECT_21A with AtEC1.2 promoter-driven expression of SpCas9. Arrows indicate promoters, Т-signs indicate terminators. SmR, KanR – spectinomycin and kanamycin resistance genes respectively. Ori pMB1 and oriV – indicate origins of replication needed for plasmid propagation in *E. coli* and Agrobacteria, respectively. LB – left T-DNA border. RB – right T-DNA border. The figure was drawn using glyphs from the SBOLCanvas web-based application (Terry et al. 2021)

For homoeolog editing, we used a modified pDIRECT_21A binary vector from Čermák et al. plant plasmid kit (Čermák et al. 2017). The 35S promoter of *A. thaliana* codon-optimized SpCas9 was substituted with the egg-cell gene promoter sequence AtEC1.2 from pMOD_A0608 of the same plasmid kit. The AtEC1.2 promoter was amplified from pMOD_A0608 using overhang PCR with the primers At_EC1.2_F (5’-ATGATCGGCGCGCCACTTTGATAAATGTTCCTCGC-3’) and At_EC1.2_R (5’-TAGCAGCCTGCAGGTTATTCTTTCTTTTTGG-3’). The amplified product was then cloned and inserted into pDIRECT_21A via the SbfI and AscI restriction sites. The egg cell-specific promoter allows the expression of the nuclease at very early stages of plant development, when the embryo consists of one or two cells, which helps to avoid chimeras in the resulting mutants (Wang et al. 2015). To test editing we have chosen two pairs of genes – orthologs of *A. thaliana SHORT VEGETATIVE PHASE (SVP)* and *CONSTANS (CO)* genes – the regulators of flowering time. Knock out phenotype of these genes is easy to identify, they have high sequence similarity (see Results section) that provides robust test for specificity of editing and are unexpected to result in lethal phenotype, even if both copies are edited.

Spacers selected for homoeolog editing (named CO_O4, CO_R1, SVP_O1, and SVP_R1; for details see Table S1) were cloned and inserted into a modified pDIRECT_21A binary vector via the GoldenGate assembly method, following protocol A of the Čermák et al. plant plasmid kit (Čermák et al. 2017) (Figure 1b).

### 3.5 Hygromycin sensitivity test

The effective concentration of hygromycin B (Invitrogen) was determined by germinating *C. bursa-pastoris* seeds of the PGL0001 accession on Petri dishes with varying concentrations of antibiotics added to the media. Prior to sowing, all the seeds were surface sterilized with a solution of 80% ethanol and hydrogen peroxide on filter paper for 5 min until the alcohol evaporated and the seeds dried. Fifty seeds were evenly distributed on MS0 media supplemented with 5 g/L Agargel (Sigma‒ Aldrich) and different concentrations of hygromycin B. The concentrations tested were 0, 10, 20, 30, 40, 50, 60, 70, 80, 90 and 100 mg/L hygromycin B per dish. Dishes with seeds were then stratified in a refrigerator at 4°C for 2 weeks, transferred to a growth room and incubated at 22°C, 50–60% humidity, and a 16/8-h day/night light cycle for another 2 weeks. The effective concentration of hygromycin B was determined as the concentration that inhibited the germination and growth of more than 90% of the seeds.

### 3.6 Bacterial strains and floral dip transformation

The *Escherichia coli* strain XL1-Blue (Evrogen, Russia) was used for routine cloning and plasmid multiplication. *Agrobacterium tumefaciens* strains GV3101 (pMP90) (Koncz and Schell 1986) and EHA105 (Hood et al. 1993) were used for plant transformation. Bacteria were transformed with plasmids via electroporation using Bio-Rad (USA) cuvettes either on “Ec1” (for *E. coli)* or “Agr” program (for Agrobacteria) of the electroporator (BioRad MicroPulser, USA) according to the manufacturer’s protocol. Transformed cells were germinated in the shaker-incubator: *E. coli –* for 1 h at 200 rpm and 37°C; *A. tumefaciens* – for 2 h at 200 rpm and 28°C. Then cells were plated on a selective LB agar medium containing an appropriate antibiotic. *E. coli* colonies containing modified pDIRECT_21A_EC1.2 vector were selected on kanamycin (50 µg/mL). *A. tumefaciens* colonies were selected on medium containing kanamycin (50 µg/mL), gentamycin (25 µg/mL) and rifampicin (25 µg/mL). For the pDOMC11 vector, the antibiotic set was the same except that spectinomycin (50 µg/mL) was used instead of kanamycin. Plasmid purification from *E.coli* was performed using the Plasmid Miniprep kit (Evrogen, Russia) according to the manufacturer’s protocol.

Two days prior to *C. bursa-pastoris’* floral dip transformation, a single colony of *A. tumefaciens,* harbouring either pDIRECT_21A_EC1.2 or pDOMC11 vector, was inoculated into 5 mL of YEB medium containing 50 mg/L rifampicin and 25 mg/L gentamicin for GV3101 (pMP90) or only 50 mg/L rifampicin for EHA105. Additionally, 50 mg/L spectinomycin or 50 mg/L kanamycin was added for agrobacteria carrying the pDOMC11 or modified pDIRECT_21A_EC1.2 binary vector, respectively. Agrobacteria were cultured overnight at 28°C in an incubator shaker at 200 rpm. The overnight culture was then diluted in 500 mL (1:100) of fresh YEB with the same set of antibiotics and grown for 18–24 h at 28°C in an incubator shaker at 200 rpm. The *A. tumefaciens* cells were then pelleted via centrifugation at 4000 g for 15 minutes at room temperature and resuspended in infiltration buffer at an OD600 of 0.8. The infiltration buffer used contained the minimum number of components required for successful transformation by floral dip of *A. thaliana* and *C. bursa-pastoris* according to published data (Clough and Bent 1998; Bartholmes et al. 2008): 5% sucrose and 0.02% Silwet-77. We also tried other compositions of infiltration medium and optical density of Agrobacterium suspension but they did not result in an increased transformation efficiencies so we used the parameters from above-mentioned studies.

Plants at an early stage of flowering (when the buds are already formed on the stem, but have not yet opened, or after the opening of the first flower) were used. Open flowers and siliques were cut off (if any), and each plant was immersed twice in the *Agrobacterium* suspension (first, the primary flower stalk and then, 7 days later, the secondary stalks from rosette axillary buds). Plants were inverted and immersed in agrobacterial suspension for 10 sec. Additionally, we dripped the suspension with a Pasteur pipette on secondary inflorescences closer to the rosette if they were problematic to dip due to the length of the stalk. Dipped plants then were covered with plastic dome to maintain high humidity and placed in the dark for 24 h. For the comparison of *A. tumefaciens* strains, a simplified version of the experiment was performed. While primary and secondary rosette flower stalks were dipped for the comparison of the six accessions, only the top 3 inflorescences of the primary flower stalk of the PGL0025 accession plants were dipped. Three pots were used per accession with 1-6 plants each (with the exception of the PGL0025 accession, which had two pots) for comparison of the six accessions, and three pots with two PGL0025 plants each for the *Agrobacterium* strain comparison were used. Seeds were collected only from the *Agrobacterium*-treated inflorescences after the siliques had matured.

### 3.7 Transgenic plant selection and transformation efficiency analysis

Given that there are no published data on the sensitivity of *C. bursa-pastoris* to hygromycin B and genetic editing vector incorporates a hygromycin resistance gene, we conducted a test to determine the effective concentration of hygromycin B for selecting transgenic *C. bursa-pastoris* seeds. The test revealed that the effective concentration for *C. bursa-pastoris* falls within the range of 20-30 mg/L (see Additional file 1, Figure S1), which is comparable to the concentrations found for *A. thaliana* selection (Ee et al. 2014). Thus, we used 25 mg/L hygromycin B (Invitrogen) for selection. The medium for selection was ½ MS1.5 supplemented with 5 g/L Agargel (Sigma‒Aldrich). Prior to sowing, all the seeds were surface sterilized with a solution of 80% ethanol and 3% hydrogen peroxide on filter paper for 5 minutes until the alcohol evaporated and the seeds dried. The Petri dishes containing sowed seeds were wrapped in foil and stratified in the dark at 4°C for one week. The samples were subsequently placed under white light at 22°C for six hours, wrapped in foil again and incubated in the dark at 22°C for two days. After incubation, the germinated seeds were screened for hygromycin-resistant seedlings that were growing vertically with elongated hypocotyls. The selected seedlings were then planted in moist vermiculite pots and grown in the growth room as described above.

For transformation efficiency studies, photographs of seeds were taken using a Zeiss Semi 508 stereomicroscope and semiautomatically counted using the Fiji software (Schindelin et al. 2012). The seeds were then screened for fluorescence using a Levenhuk MED PRO 600 Fluo microscope excited by green light (510-548 nm) and observed through a red cutoff filter (585-700 nm). Photographs of the transgenic fluorescent seeds were obtained using ToupView software. The transformation efficiency was determined by calculating the percentage of red fluorescent seeds (tagRFP+) among the samples of seeds collected and analyzed per pot for the comparison of six accessions and per plant for *Agrobacterium* strain comparison (for details see Table S3 and S4).

To compare transformation efficiency across six accessions, we calculated the ratio of transformed seeds to total seeds sampled per pot. The total number of seeds analyzed per pot ranged from 1,749 to 6,369, with a grand total of 86,406 seeds analyzed. Each accession was evaluated using three independent replicates (pots), except for PGL0025, which had two replicates, resulting in a total sample size of n = 17. Normality assumptions were confirmed for all groups using Shapiro-Wilk tests (p > 0.05). To account for variability in plant density between pots, we used Spearman’s rank correlation to assess potential confounding effects. The correlation analysis revealed no significant association between transformation efficiency and plant density (p = 0.228), justifying the exclusion of plant density as a covariate. Homogeneity of variances was confirmed by the Brown-Forsythe test (p = 0.260), satisfying ANOVA assumptions. Consequently, accession comparisons were performed using one-way ANOVA, followed by Tukey’s post-hoc test for pairwise differences.

To compare Agrobacterium strains for transformation efficiency, we calculated the ratio of transformed seeds to total seeds sampled per pot. The total number of seeds analyzed per pot ranged from 991 to 3,395, with a grand total of 11,490 seeds analyzed. Data were aggregated at the pot level to align with the experimental design and avoid pseudoreplication. Each strain had three replicates (individual pots treated as independent replicates), yielding a total sample size of n = 6. Normality was assessed using Shapiro-Wilk tests, which confirmed that the data for GV3101 met the normality assumptions (p > 0.05), whereas those for EHA105 did not. Thus, we conducted both parametric Welch’s t-test and nonparametric Mann-Whitney U-test analyses to assess differences in efficiency between the strains.

## 4 Results

### 4.1 PGL0001 gRNA database development and the potential of differential editing of homoeologs

To provide genome-wide characterization of the possibility of homoeolog-specific genetic editing, we developed a database that contains comprehensive information on the location and characteristics of all spacers with the canonical SpCas9 “NGG” PAM motif in the PGL0001 genome (see Materials and methods for details). The filtering process left 2,487,380 spacers in the database. These spacers are located in the CDS of 60,468 genes, representing 94.8% of the total number of annotated genes (63,768).

A total of 22,776 1-to-1 homoeolog pairs were identified in the genome of *C. bursa-pastoris* using OrthoFinder software (Emms and Kelly 2015, 2019). To investigate the potential for differential editing of these genes, the gRNAs corresponding to these genes were extracted from the abovementioned database. Furthermore, those gRNAs that have potential off-targets with CFD ≥ 0.05 (≥ 5% potential off-target activity) were excluded, leaving 26,531 spacers specific to the homoeologs.

As demonstrated by the metric values of the selected homoeolog targeted gRNA, these spacers are likely to function effectively. The first quartile of scores for DeepSpCas9 is above 0.36, whereas for DeepHF, it is above 0.5. This finding indicates that the majority of gRNAs from the selected set should have a predicted on-target editing efficiency of 36% or more. The first quartile of the aggregated CFD score is above 0.95, indicating that the majority of spacers should demonstrate minimal off-target activity. However, while there are outliers with scores below 0.9, these belong to the gRNA with a high number of off-targets with CFD < 0.05, which decreases the aggregate score, but should not affect specificity (Figure 2a).

**Fig.2.**
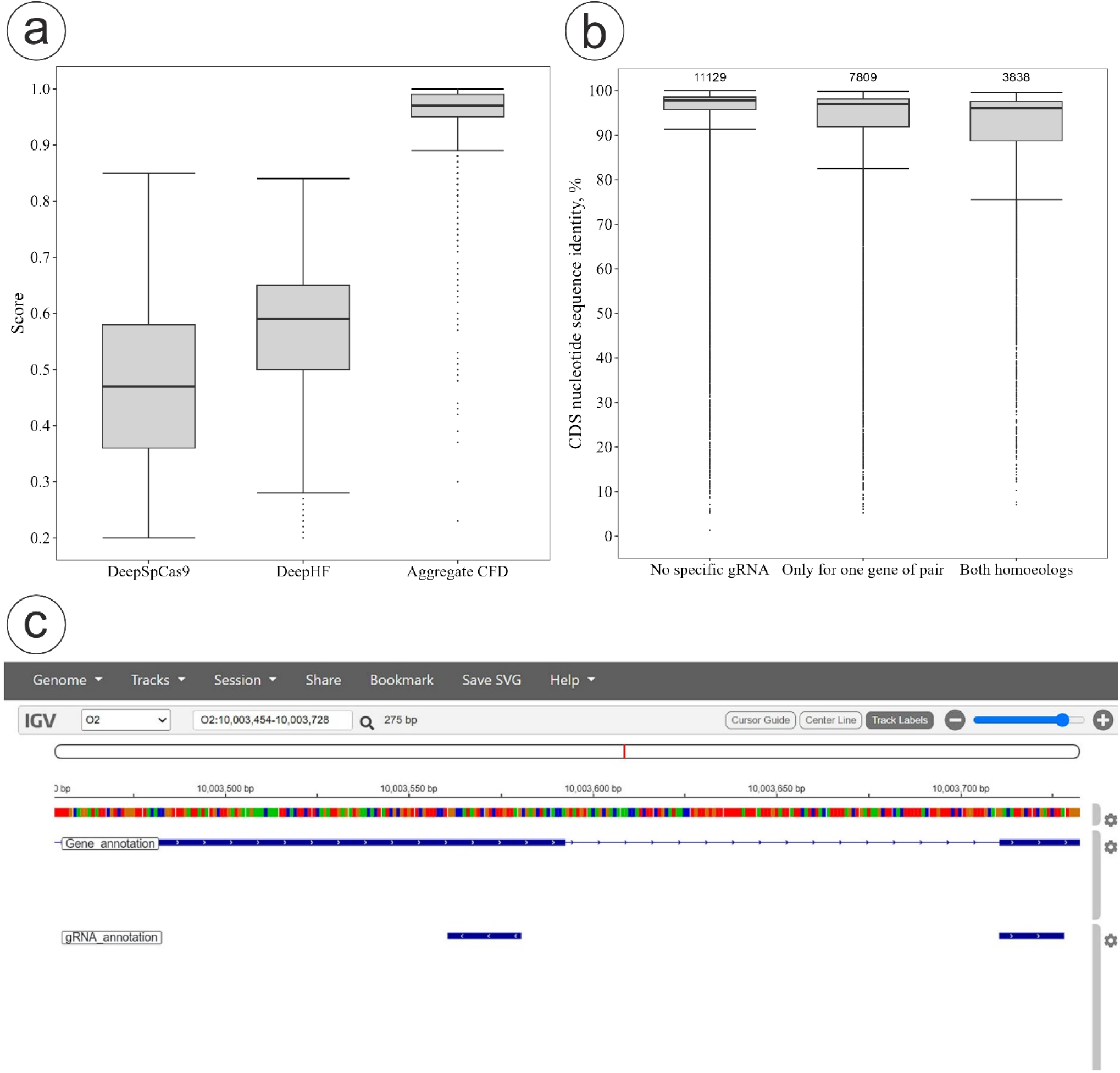
(a) Boxplot of gRNA on- and off-target metrics, (b) boxplot of the range of CDS nucleotide similarity of homoeolog pairs with differential editing capability, numbers above the boxes represent the number of homoeolog pairs that fall into each category, (c) the view of *the C. bursa–pastoris* genome browser showing gene annotation and gRNA position. Panels (a) and (b) were constructed using the R library ggplot2 v3.4.0 (Villanueva and Chen 2019)

To determine whether there is a relationship between the nucleotide sequence similarity of homoeolog pairs and the ability of gRNAs for differential editing, we classified pairs of homoeologs in the filtered database into three groups: pairs for which no specific gRNAs were found for both genes in pair, pairs for which specific gRNAs were found only for one homoeolog in the pair, and pairs for which specific gRNAs were identified for both homoeologs. The results revealed that no specific gRNAs for differential editing could be identified for 48.9% of the homoeolog pairs for *C. bursa-pastoris*. In 34.3% of the pairs for *C. bursa-pastoris* only one homoeolog of the pair could be specifically edited, whereas in 16.9% of the pairs for *C. bursa-pastoris* both homoeologs could be differentially edited. As anticipated, the majority of pairs devoid of suitable gRNA for differential editing exhibited high degrees of sequence similarity, exceeding 95%. However, no clear distinction could be identified between the groups, and the ranges of sequence similarity exhibited significant overlap (Figure 2b). These results demonstrate that reliable prediction of differential editing potential in homoeologous gene pairs based solely on sequence similarity is not feasible.

To make the results of the genome-wide selection of gRNAs available for the community, we have published database as well as list of identified homoeolog pairs on Figshare (Omelchenko et al. 2024a), and we have integrated gRNA data into the publicly available IGV genome browser (https://t2e.online/igv_capsella_bursa-pastoris/). It shows the position of gRNAs along with the annotation of protein-coding genes (Figure 2c); by clicking on a certain gRNA, the user can see the information on the position of the gRNA, its metrics, the targeted locus and other features.

### 4.2 Guide RNA selection for differential editing of homoeologous genes

*CO* and *SVP* genes in *C. bursa-pastoris* demonstrate high pairwise identity of coding sequences between subgenomes. The aligned *SVP* coding sequences have 11 SNPs and an indel of 15 nucleotides with a pairwise identity of 98.41% (indels not counted), and the *CO* coding sequences have 36 SNPs and four indels of 36, 3, 111, and 24 nucleotides in length, respectively, with a pairwise identity of 96.7% (indels not counted) (alignments are presented in Additional file 1, Figures S2 and S3). Thus, the homoeologs of the *SVP* gene exhibit a high degree of similarity, which may pose challenges in finding gRNAs for differential editing. Conversely, the homoeologs of the *CO* gene display a lower degree of similarity and contain large indels, thereby increasing the likelihood of identifying specific spacers.

We selected gRNA for each homoeolog of the *CO* and *SVP* genes from the top 10 gRNAs ranked in the full database based on their sequence characteristics and predicted on- and off-target activity using the crisprVerse in-built ranking function (selected gRNA details are presented in Additional file 2, Table S2). All selected spacers are located on the first exon close to the start codon of the *CO* and *SVP* homoeologs (see Additional file 1, Figure S4).

### 4.3 SpCas9:sgRNA *in vitro* cleavage assay

Prior to *in vivo* experiments, we explored the efficiency and specificity of editing using *in vitro* cleavage assay. DNA fragment analysis of the CO_O and CO_R amplicons after *in vitro* reactions with SpCas9 and sgRNA demonstrated efficient cleavage of their respective target homoeologs for both sgRNAs, resulting in fragments of the expected length (Figure 3a, b). However, CO_R1 sgRNA also induced off-target cleavage of CO_O despite the noncanonical PAM motif “CCG” in CO_O and the absence of CO_O among the predicted off-targets in the database, although with low efficiency (the major peak is the intact CO_O amplicon, Figure 3a). The analysis of SVP_O and SVP_R amplicons after *in vitro* reactions with SpCas9 and sgRNA also demonstrated efficient cleavage of their respective target homoeologs for both sgRNAs (Figure 3c, d). However, in contrast to the results for *CO* homoeologs, both sgRNAs cleaved their nontarget homoeologs as well, despite one mismatch in each sgRNA between the spacer sequences in the seed region of the spacer and the corresponding protospacer of the nontarget homoeolog. However, SVP_O1 showed reduced off-target cleavage efficiency, as evidenced by the almost equal height of the intact SVP_R amplicon peak and cleaved fragments (Figure 3d). Thus, on the basis of *in vitro* assays, none of the selected sgRNAs are able to specifically edit their corresponding SVP homoeologs.

**Fig.3.**
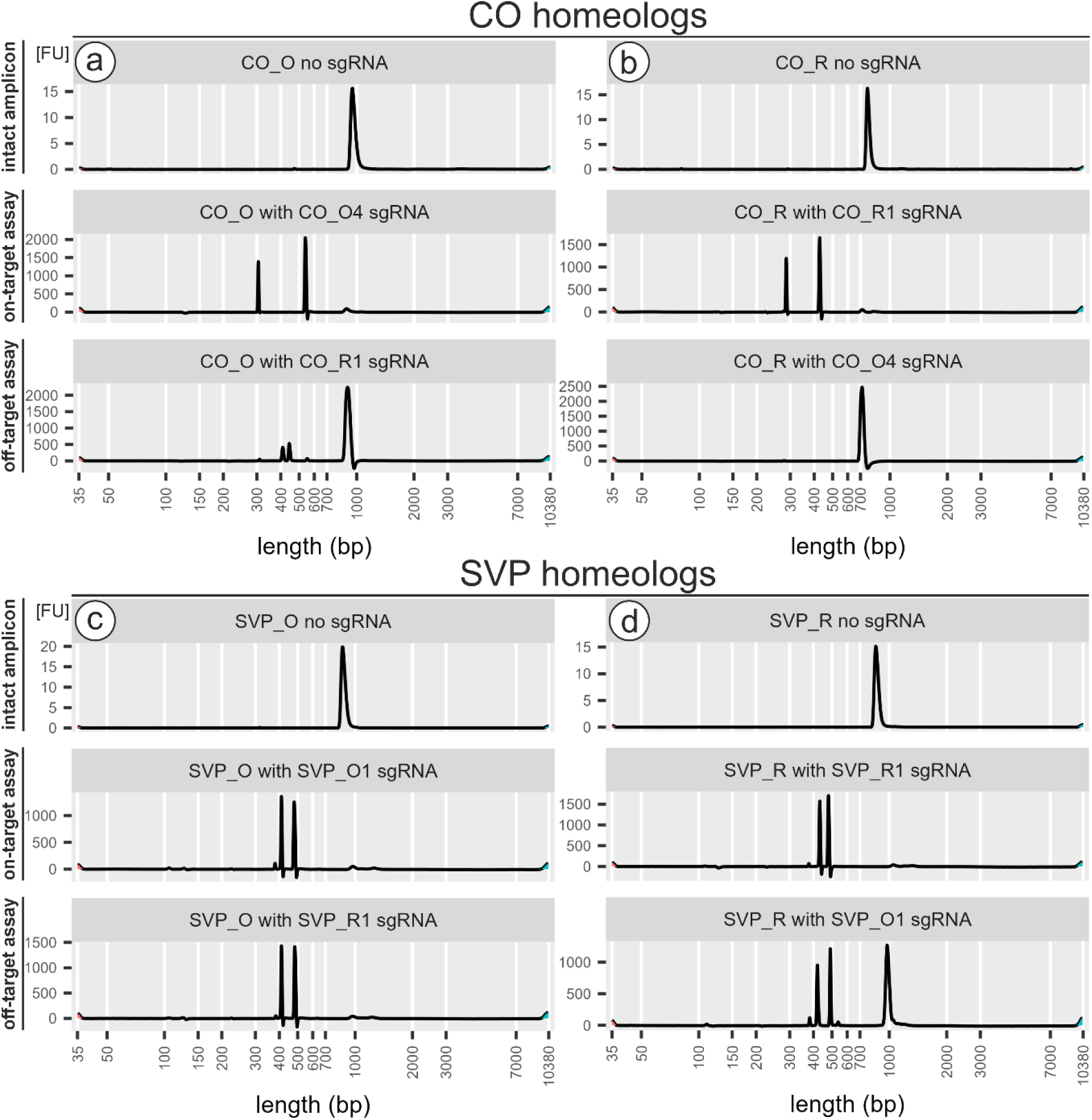
Electrophoregrams of *CO* and *SVP* homoeolog amplicons after *in vitro* cleavage assay with SpCas9 and sgRNA. (a) CO_O intact amplicon, on-target test with CO_O4 sgRNA, and off-target test with CO_R1 sgRNA; (b) CO_R intact amplicon, on-target test with CO_R1 sgRNA, and off-target test with CO_O4 sgRNA; (c) SVP_O intact amplicon, on-target test with SVP_O1 sgRNA, and off-target test with SVP_R1 sgRNA; (d) SVP_R intact amplicon, on-target test with SVP_R1 sgRNA, and off-target test with SVP_O1 sgRNA. Plots were generated in R using bioanalyzeR package v0.10.1 (Foley 2024)

### 4.4 *CO* and *SVP* homoeolog-aware editing

The transformation of *C. bursa-pastoris* accession PGL0001 with *A. tumefaciens* GV3101 (pMP90), which carry vectors with CRISPR/Cas and gRNA expression cassettes, and further selection on hygromycin-containing media resulted in the isolation of T1 transgenic plants. For CO_O sgRNA, two T1 plants were selected; however, Sanger sequencing revealed that only one of them had the mutation. This plant had a homozygous “A” insertion 3 nucleotides upstream of the PAM in the *CO_O* homoeolog, causing a frameshift mutation (Figure 4a), whereas the *CO_R* homoeolog remained intact (Figure 4b). In the T1 generation grown from seeds of plants transformed with the CO_R sgRNA construct, 3 out of 15 transgenic plants presented a heterozygous mutation in the *CO_R* homoeolog. To obtain homozygous mutants in the T2 generation, we planted 40 seeds from T1 mutant plants. In 10 of the grown plants, we detected a homozygous frameshift mutation in the target gene due to a “C” nucleotide insertion (Figure 4c), whereas the *CO_O* homoeolog remained intact (Figure 4d).

**Fig.4.**
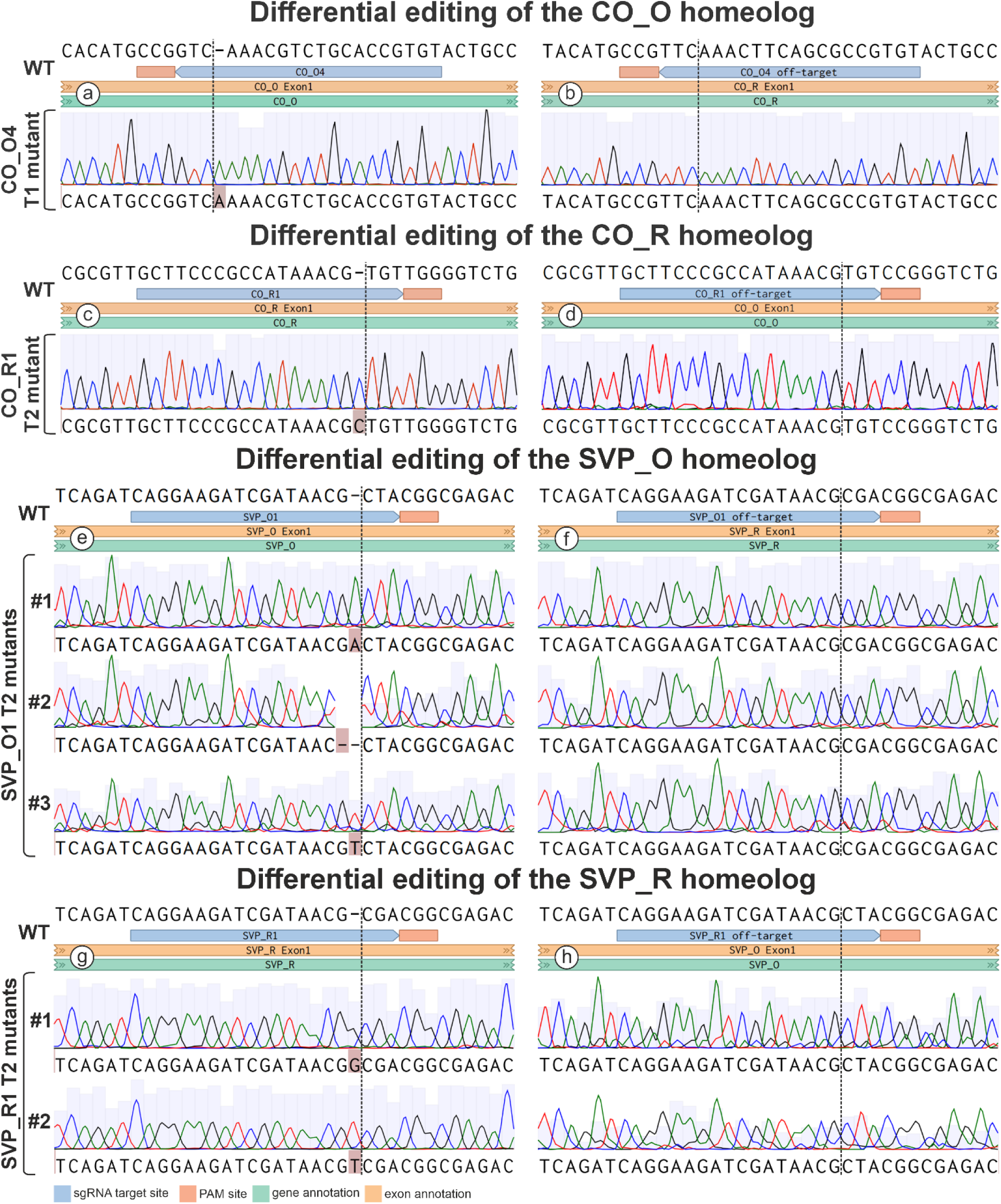
Sanger sequence alignments of wild-type (WT) plant compared with T1 *co_o*, T2 *co_r,* T2 *svp_o* and T2 *svp_r* mutant plants. Alignment of CO_O4 sgRNA on-target regions in *CO_O* (a) and off-target regions in *CO_R* (b) homoeologs; alignment of CO_R1 sgRNA on-target regions in *CO_R* (c) and off-target regions in *CO_O* (d) homoeologs. Alignment of SVP_O1 sgRNA on-target region in *SVP_O* (e) and off-target region in *SVP_R* (f) homoeologs; alignment of SVP_R1 sgRNA on-target region in *SVP_R* (g) and off-target region in *SVP_O* (h) homoeologs. The indels are highlighted in pink. Figures were created from screenshots of MAFFT alignments in the Benchling web-service (https://www.benchling.com/)

For SVP_O sgRNA transformation, only one T1 plant, which had a heterozygous mutation in the *SVP_O* homoeolog, was obtained. Three out of 20 T2 plants grown from seeds of each had a distinct homozygous frameshift mutation: an “A” insertion, a “T” insertion, and a 2 bp deletion in the cleavage region of SVP_O1 (Figure 4e). The *SVP_R* homoeolog had no mutations in any of the T2 plants, contrary to expectations based on an *in vitro* assay (Figure 4f). For SVP_R sgRNA, two T1 transgenic plants with a heterozygous mutation were obtained. Among 40 T2 plants, 19 plants presented a homozygous frameshift mutation in the *SVP_R* homoeolog. Two of these plants had a “G” insertion, whereas the remaining 17 plants had a “T” insertion (Figure 4g). The *SVP_O* homoeolog remained intact in all T2 plants (Figure 4h).

Thus, despite the results of the computational estimation of gRNA specificity and the *in vitro* SpCas9:sgRNA assay, we obtained plants with specific differential editing for each of the homoeologs of the *CO* and *SVP* genes. This highlights the incongruence of the outcomes of *in vitro* and *in vivo* editing, with the latter being more specific.

### 4.5 *Agrobacterium* transformation efficiency in *C. bursa-pastoris* accessions

We compared efficiency of *Agrobacterium*-mediated transformation for six accession of *C. bursa-pastoris* representing geographically and phylogenetically distant populations. The screening of the seeds from the transformed plants revealed that all the accessions contained transgenic seeds with varying levels of fluorescence intensity (see Figure 5a, b). The transformation efficiency was highest in the accession PGL0025 (‘KBG’, Asian clade), with a rate of 1.07-1.09%. This value is 4.7 times greater than that of the second most efficiently transformed apetalous accession, PGL0002. The Murmansk accession PGL0066 completes the top three, while the other three accessions showed low transformation efficiency of less than 0.1% (for details see Additional file 2, Table S3). A one-way ANOVA revealed a significant difference among the group means (p < 0.0001). Tukey’s multiple comparisons test further demonstrated significant differences between the transformation efficiencies of accessions (Figure 5c).

**Fig.5.**
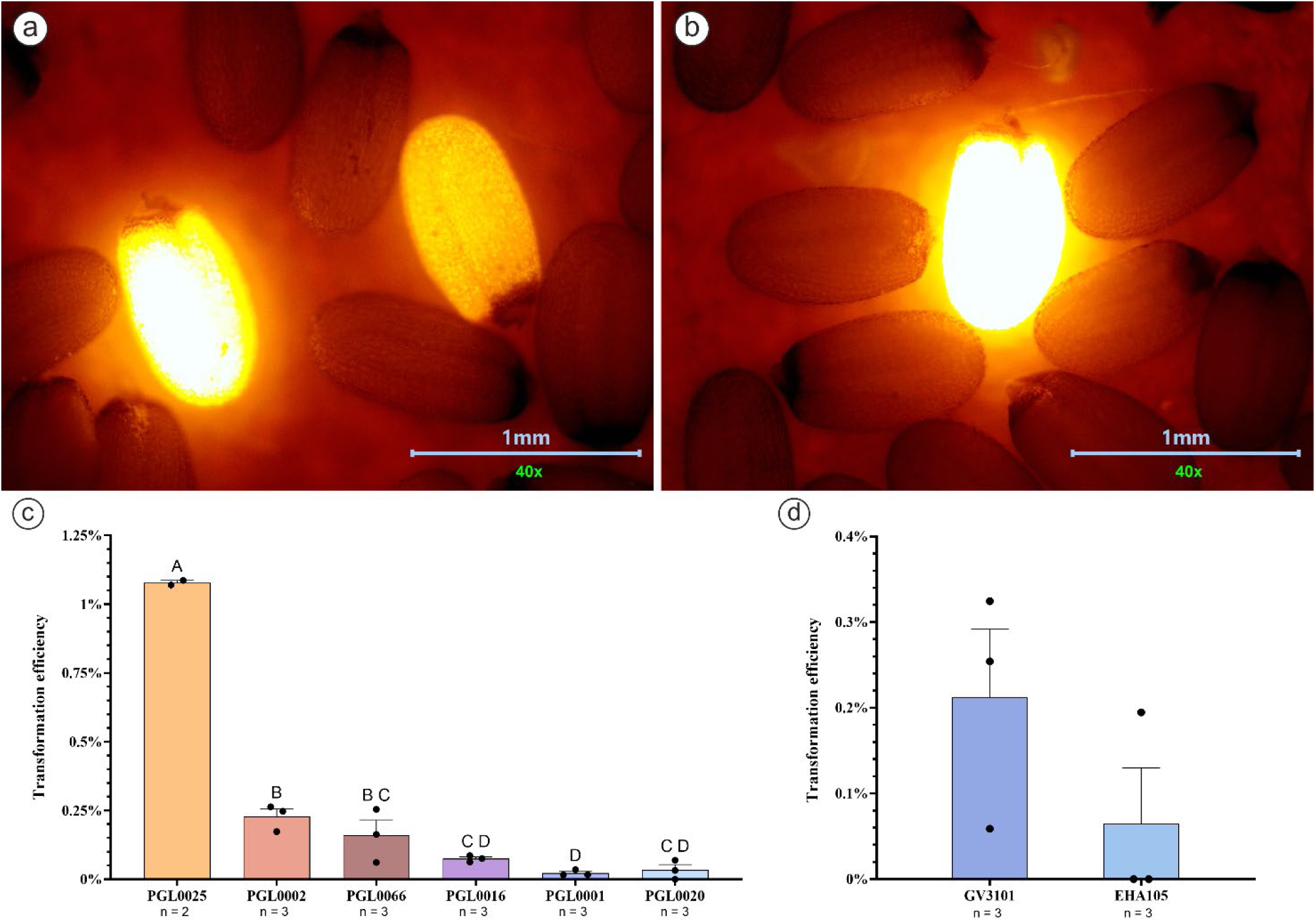
Examples of bright and dim (a) and very bright (b) tagRFP+ seeds of *С. bursa-pastoris* PGL0025 accession; (c) Transformation efficiency as a percentage of tagRFP+ seeds/collected seeds per pot. Means not sharing any letter are significantly different by the Tukey’s test at α = 0.05, dots depict individual values; (d) GV3101 (pMP90) versus EHA105 transformation efficiency of the PGL0025 accession counted as a percentage of tagRFP+ seeds/collected seeds per pot; dots depict individual values. Figures (c) and (d) were generated using GraphPad Prism software, for both figures error bars represent Mean ± SEM.

In addition, we compared the transformation efficiency of the PGL0025 accession between the *A. tumefaciens* strains GV3101 and EHA105 in a reduced version of the experiment (see Materials and methods). The number of tagRFP+ seeds indicated that GV3101 (pMP90) was more effective at transforming *C. bursa-pastoris* PGL0025 accession (Figure 5d). Five out of the six plants transformed with GV3101 (pMP90) produced fluorescent seeds, whereas only two out of the six plants transformed with EHA105 produced such seeds (for details see Additional file 2, Table S4).

Statistical comparison of transformation efficiency between Agrobacterium strains revealed no significant differences. Welch’s t-test showed comparable efficiencies (t=1.439, p = 0.2262) between GV3101 (Mean ± standard error of the mean (SEM): 0.21% ± 0.08%) and EHA105 (0.06% ± 0.07%). Similarly, the non-parametric Mann-Whitney U-test corroborated this finding (U = 1.0, p = 0.2000). While statistical significance may be marginal due to limited replicates, the effect size suggests GV3101 is superior.

## 5 Discussion

*Capsella bursa-pastoris* and the whole genus *Capsella* are long-standing models for many areas of plant biology (for review, see (Sicard and Lenhard 2018)), yielding discoveries in developmental, population and ecological genetics (e.g., (Nutt et al. 2006; Slotte et al. 2007; Kryvokhyzha et al. 2016)). However, functional genetics studies on this species are lagging behind. Our study, which uses a recently assembled genome sequence and a set of geographically distant accessions, fills this gap in two aspects: 1) the development of a gRNA database with a focus on homoeolog-specific editing and 2) the selection of an accession with a high transformation rate.

Genetic editing has become a tool of choice for both fundamental and applied research in functional genetics. Databases of gRNAs and concomitant tools (search engines, genome browsers) are essential to fully unlock its potential. Such databases were developed for model and agricultural species (Xie et al. 2014; Park et al. 2016; Cram et al. 2019). An important condition for the successful design of genetic editing experiments is the availability of genome sequences. Until recently, two assemblies were published for C. *bursa-pastoris* – in 2015 (Douglas et al. 2015) and in 2017 (Kasianov et al. 2017). In the first one, only 40% of the assembly was divided into homoelogous sequences; for the second, this fraction is much higher but the number of homoeologous pairs is still quite low (∼ 16 thousand pairs), and the genome is not chromosome-scale; thus, the assignment of genes to subgenomes is not conclusive. Using a recently published chromosome-scale genome sequence with better resolution of homoeologs (∼ 23 thousand pairs), we performed genome-wide development of the gRNA database and represented it in a publicly available form in the genome browser. An *in silico* search of gRNAs suggested that only a small number of homoeologs can be specifically edited.

We complemented this *in silico*-based part of the study by an experimental test and optimization on a model polyploid *C. bursa-pastoris*. As mentioned above, an *in silico* search of gRNAs suggested that only a small number of homoeologs can be specifically edited. We performed an experimental check (both *in vitro* and *in vivo*) of the possibility of specific editing for two pairs of homoeologs. The results of the *in vitro* cleavage assay showed that it tends to overestimate non-target activity as it revealed off-target activity leading to the cutting of nontarget homoeologs while *in vivo* experiments revealed that this does not occur. This finding is congruent with the findings of Fu and coauthors (Fu et al. 2016), who showed that SpCas9 is more tolerant to mismatches in the spacer sequence *in vitro* than it is *in vivo*. In plants, few studies address this question, but ones that do (Sagarbarria et al. 2023) yield similar conclusions. Based on these results, *in vitro* cleavage test is the most useful for revealing poorly performing gRNAs (ones that do not perform well *in vitro* will not perform better *in planta* - but the opposite is not necessarily true).

Another important component of the functional genetics toolkit is the transformation. This method allows the generation of plants with heterologous genes, altered gene expression levels and/or patterns, and the integration of reporter constructs. In the context of genetic editing, introducing the expression of the CRISPR associated editing enzyme (usually Cas9) and a gRNA serves as the first step. Owing to its simplicity and speed, the floral dip method of *Agrobacterium*-mediated transformation has become the method of choice for *A. thaliana* and related plants. However, it is important to note that the efficiency of floral dip transformation is dependent on the plant’s genotype, as various ecotypes/accessions/cultivars are transformed with different efficiencies. For example, *A. thaliana* ecotypes Ws-O, Nd-O, No-O, and Col-O are transformed with an efficiency of 0.5-3%, whereas ecotypes Ler-O, Dijon-G, and Bla-2 are transformed with an efficiency 10-100 fold lower (Clough and Bent 1998). A dependence of transformation efficiency on plant genotype was also observed for *C. bursa-pastoris* in an earlier study (Bartholmes et al. 2008). The authors were able to achieve transformation efficiencies of up to 0.4% for the ‘wt 6/1+2’ line but not for the ‘Spe 9/9’ and ‘Spe 2/4’ lines under the same experimental conditions. In this study, we compared the transformation efficiency of six *C. bursa-pastoris* accessions - wild type and apetalous mutant from Moscow (Russia), as well as from populations collected in Murmansk (Russia), Tbilisi (Georgia), Kunming (China), and Jerusalem (Israel), which represent distant clades within the highly polymorphic species *C. bursa-pastoris* (Penin et al. 2024) and observed clear differences. The most efficiently transformed accession is the PGL0025 of *C. bursa-pastoris* from China, with a transformation efficiency of ∼1%, which is higher than previously published data on the efficiency of floral dip *C. bursa-pastoris* and comparable to the transformation efficiency of a model ecotype of *A. thaliana*, Col-0. Notably, the PGL0025 ecotype stands out for its rapid flowering and compact size, offering significant advantages in terms of convenience for floral dip transformation, time and labor efficiency compared to other accessions. These characteristics make PGL0025 a highly promising candidate for future research involving transgenic *C. bursa-pastoris*.

The choice of *Agrobacterium* strain is also important factor affecting transformation efficiency. We compared two strains - GV3101 (pMP90) and EHA105. Our results indicate that GV3101 (pMP90) is superior in transforming *C. bursa-pastoris*. This finding is congruent with the results of Bartholmes et al., who compared the transformation efficiencies of LBA4404 and GV3101 (pMP90), and found latter more effective. The study on *A. thaliana* (Ghedira et al. 2013) comparing the transformation ability of GV3101 (pMP90) and LBA4404 also showed the superiority of the former.

## 6 Conclusions

Our results provide researchers working on *C. bursa-pastoris* with genome-wide information on gRNAs targeting protein-coding genes and show that selective editing of homoeologs is possible even in the case of very high similarity, highlighting that both *in silico* prediction and *in vitro* cleavage test overestimate the probability of non-specific editing. Also, we compared the transformation efficiencies for six *C. bursa-pastoris* accessions and found one that has the transformation rate as high as 1%. We expect that these findings and resources will promote polyploidy studies that use this plant as a model in functional genetics experiments.

## 7 Statements and Declarations

### 7.1 Ethics approval and consent to participate

Not applicable

### 7.2 Consent for publication

Not applicable

### 7.3 Data availability statement

Sanger sequencing results of *CO_O*, *CO_R*, *SVP_O*, and *SVP_R* amplicons of *C. bursa-pastoris* mutant plants to check on-target and off-target results are available from Figshare (Omelchenko et al. 2024b). The sequencing results supporting the findings can be accessed at GenBank under the following accession numbers: PP711553-PP711570. The *C. bursa-pastoris* gRNA database as a TSV file, as well as list of identified homoeolog gene pairs, are available from Figshare (Omelchenko et al. 2024a). Other materials (e.g. seeds, vectors) and data used and/or analyzed during the current study are available from the corresponding author upon request.

### 7.6 Authors’ contributions

ODO and LMD conceived and designed the experiments and wrote and revised the manuscript; ODO, BAM and OLV grew the plants, cultured the bacteria and performed the *Agrobacterium*-mediated transformations; ODO and BAM constructed the binary vectors, performed the *in vitro* cleavage assay, analyzed and validated the results of the experiments; ODO created the sgRNA spacer database, performed the hygromycin sensitivity test, made figures for the manuscript; BAM screened the plants for mutations in the homoeolog-aware editing experiments; and KAV and PAA established the original collection of the Shepherd’s purse accessions and provided seeds for experimentation.

All the authors read and approved the final manuscript.

## 7.7 Acknowledgements

The authors would like to thank Pozdyshev D.V. (Lomonosov Moscow State University, Moscow, Russia) for GV3101 (pMP90), Ivanova A.S. (Vavilov Institute of General Genetics, Moscow, Russia) for EHA105 *A. tumefaciens* strains, Schelkunov M.I. (Institute for Information Transmission Problems of the Russian Academy of Sciences, Moscow, Russia) for integrating sgRNA track to *Capsella* genome browser.

## 8 Additional files

### Additional file 1.pdf

**Fig.S1** *C. bursa-pastoris* PGL0001 seed germination after 2 weeks with different concentrations of hygromycin B

**Fig.S2** The alignment of the CDS of homoeologs of the CO gene. Differences between sequences are highlighted in red. The figure was created using the Benchling web service (https://www.benchling.com/)

**Fig.S3** The alignment of the CDS of homoeologs of the SVP gene. Differences between sequences are highlighted in red. The figure was created using the Benchling web service (https://www.benchling.com/)

**Fig.S4** Locations of the selected spacers in the first exons of *CO* (a) and *SVP* (b) homoeologs. For each homoeolog sgRNAs used in the study are indicated with blue color below the amino acid sequences, adjacent PAM sites are marked with orange. Below gRNA annotations 1st exons of CO and SVP genes are marked with yellow, and whole gene marked with light green, red highlighted nucleotides mark mismatches between homoeolog sequences. Figures were created from screenshots of MAFFT alignments made in the Benchling web-service (https://www.benchling.com/)

### Additional file 2.xlsx

**Table S1.** Characteristics of primers for amplification of fragments of the homoeologs of *CO* and *SVP* genes. The IDs’ prefixes are presented in the format ‘gene_subgenome’

**Table S2.** Selected sgRNA summary characteristics

**Table S3.** Floral dip transformation efficiency of six accessions using GV3101 (pMP90) strain

**Table S4.** Comparison of the floral dip transformation efficiency between GV3101 and EHA105 strain for the PGL0025 accession

