## Additional file 1 for "Optimization of functional genetics tools for a model tetraploid Capsella bursa-pastoris, with focus on homoeolog-aware gene editing"

**Fig.S1** *C. bursa-pastoris* PGL0001 seed germination after 2 weeks with different concentrations of hygromycin B

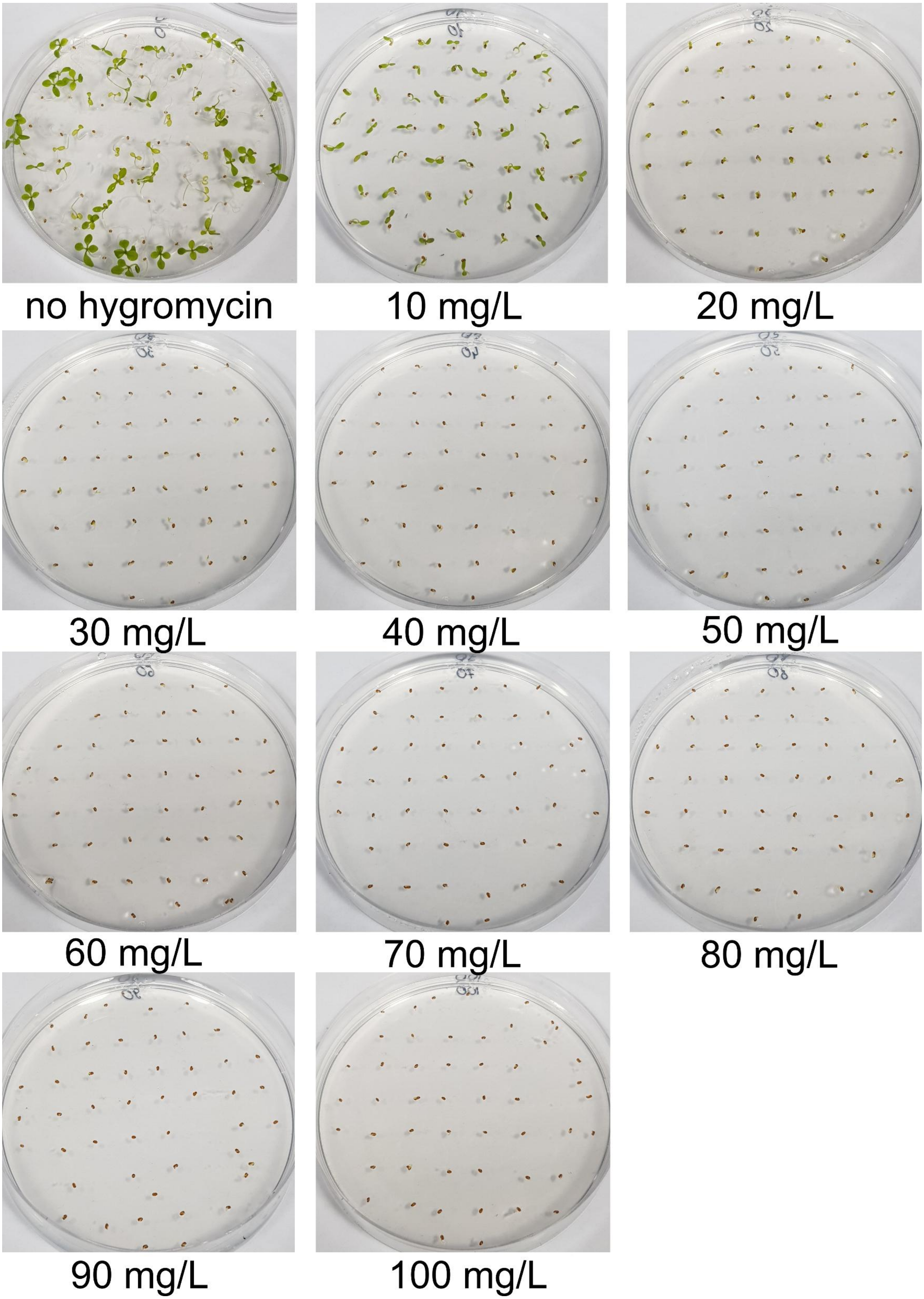

**Fig.S2** The alignment of the CDS of homoeologs of the CO gene. Differences between sequences are highlighted in red. The figure was created using the Benchling web service (https://www.benchling.com/)

```
            1                                                                              80
CO_R_msk_CDS ATGTTAATACAAGAGAGTAACGATATTAGTAGTGGAGAGAACAACACGGCACGAGCCTGTGATACATGCCGTTCAAACTT
CO_O_msk_CDS ATGTTAAAACAAGATAGTAACGATATTAGTAGTGGAGAGAACAACAGGGCACGAGCCTGTGACACATGCCGGTCAAACGT

            81                                                                            160
CO_R_msk_CDS CAGCGCCGTGTACTGCCATGCAGATTCAGCCTACTTGTGCATGAGCTGTGATGCTCAAGTTCACTCTGCCAATCGCGTTG
CO_O_msk_CDS CTGCACCGTGTACTGCCATACAGATTCAGCCTACTTGTGCATGAGCTGTGATGCTCAAGTTCACTCTGCCAATCGCGTTG

            161                                                                           240
CO_R_msk_CDS CTTCCCGCCATAAACGTGTTGGGGTCTGCGAGTCAT------------------------------------ATGATGCA
CO_O_msk_CDS CTTCCCGCCATAAACGTGTCCGGGTCTGCGAGTCATGTGAGCGTGCTCCAGCTGCTTTTTTGTGCGAGACAGATGATGCA

            241                                                                           320
CO_R_msk_CDS TCTCTATGCACAGCCTGTGATTCAGAGGTTCATTCTGCAAATCCACTTGCTAGACGCCATCACCGAGTTCCAATTCTGCC
CO_O_msk_CDS TCTCTATGCACAGCCTGTGATTCAGAGGTTCATTCTGCAAACCCACTTGCTAGACGCCATCACCGAGTTCTAATTCTGCC

            321                                                                           400
CO_R_msk_CDS AATCTCCGGAAACTCGTACAGCTCCATGGCCACTCACCACCGAAGCCAGAGAACAATGGACGATCAAGAGAAGATACCAG
CO_O_msk_CDS AATCTCCGGAAACTCGTACAGCTCCATGGCCACTCACCACCGAAGCGAGAGAACAATGAACGATCAAGAGAAGATACCAG

            401                                                                           480
CO_R_msk_CDS TGGTGGATGAAGAGACAAGTAAGGAGGCTAATGAGGATGCCAAGGAGGTGGTGTCTTGGTTGTTCCCTAACTCAGATAAA
CO_O_msk_CDS TGGTGGATGAAGAGACAAGTAAGGAGGCTAATGAGGATGCCAAGGAGGTGGTGTCTTGGTTGTCCCCTAACTCAGATAAA

            481                                                                           560
CO_R_msk_CDS AATAGTGGCAATCATAACAACAATAACCAGAACAATGTGTTATTGTTTAGTGATGCGTATACAGACCTTGCGGATCACAA
CO_O_msk_CDS AATAGTGGCAATTATAACAACAATAACCAGAACAATGTGTTATTGTTTAGTGATGCGTATCTAGACCTTGCGGATCACAA

            561                                                                           640
CO_R_msk_CDS CTCAAGTATGGACTACCAATTCACAGGTCAATACAATCAACAAGAACAGGACTGTGGTGTACCACAAACGAGCTACGGTG
CO_O_msk_CDS CTCAAGTATGGACTACCAATTCACAGGTCAATACAATCAAC---AGCAAGACTGTGGTGTACCACAAACGAGCTACGGTG

            641                                                                           720
CO_R_msk_CDS GAGATAGGGTGGTTCCACTTCAACTTGAAGAATCAAGGGGACACCTGAACCACAATGAACATAATTTTCAGTTCGATATC
CO_O_msk_CDS GAGATAGGCTGGTTCCACTTCAACTTGACGAATCAAGGGAACACCTGAACCACAATGAACATAATTTTCAGTTCGATATC

            721                                                                           800
CO_R_msk_CDS ACATATGGCTCCTCAGGAAGCCACTACACCGACAAAGGTTCTGAAAACAATAATAAATTTCACCATAAGTCTATAATAAT
CO_O_msk_CDS ACATATGGCTCCTCAGGAAGTCACTACACCGACAAAGGTTCTCAAAACAATAAT--------------------------

            801                                                                           880
CO_R_msk_CDS AACAAAGAAAAATATGCTTCCTTATGCATCGACTCTCTTTAGTCCTCTTACCAGGGGGATTGAGTGTCTTTGTTTCTGTC
CO_O_msk_CDS --------------------------------------------------------------------------------

            881                                                                           960
CO_R_msk_CDS ATTGGGCATACATTTCATCCATCGAAACTGGTGTTGTGCCGGTGTCAACAGCATGTTTCACAACAGCTTCACATCCAAGA
CO_O_msk_CDS -----GCATGCATTTCATCCATCGAAACTGGTGTTGTGCCGGTGTCAACAGCATGTTTCACAACAGCTTCACATCCCAGA

            961                                                                          1040
CO_R_msk_CDS ACTCCCAAAGCGACTATAGACAAACTACCTGACCCTCCAAGTCAGATGGTAACCCAACTCTGTCAAACGGACAGAGAAGC
CO_O_msk_CDS ACGCCCAAAGCGACAATAGACAAACTACCTGACCCTCCAAGTCAGATGGTAACCCAACTCAGTCCAACGGACAGAGAAGC

            1041                                                                         1120
CO_R_msk_CDS CAGGGTCCTGAGATACAGAGAGAAGAGGAAGACGAGGAAATTTGAGAAGACGATAAGGTATGCTTCAAGAAAAGCATATG
CO_O_msk_CDS CAGGGTCCTGAGATACAGAGAGAAGAGGAAGACAAGGAAATTTGAGAAGACGATAAGGTATGCTTCAAGAAAAGCATATG

            1121                                                                         1200
CO_R_msk_CDS CAGAGAGAAGGCCAAGGGTCAATGGTCGTTTTGCAAAGAGGATAGAAATCGAAGATGAGGACCATGGGTTCAACACAATG
CO_O_msk_CDS CAGAGAGAAGGCCAAGGGTCAATGGTCGTTTCGCAAAGAGGATAGAAGTCGAAGATGAGGACCATGGGTTCAACACAATG

            1201                                                   1266
CO_R_msk_CDS CTAATGTACGAAACAGGGTATGGCGTTGTTCCTTCATTCTG------------------------A
CO_O_msk_CDS CTAATGTACGAAACAGGATATGGCGTTGTTCCTTCATTCTGTGTTATAGGAAATATGAGGAGCTGA
```

**Fig.S3** The alignment of the CDS of homoeologs of the SVP gene. Differences between sequences are highlighted in red. The figure was created using the Benchling web service (https://www.benchling.com/)

```
                1                                                                              80
SVP_R_msk_CDS   ATGGCGAGAGAAAAGATTCAGATCAGGAAGATCGATAACGCGACGGCGAGACAAGTGACGTTCTCGAAACGAAGAAGAGG
SVP_O_msk_CDS   ATGGCGAGAGAAAAGATTCAGATCAGGAAGATCGATAACGCTACGGCGAGACAAGTTACGTTCTCGAAACGAAGAAGAGG

...............................................................................................

                81                                                                            160
SVP_R_msk_CDS   TCTTTTCAAGAAAGCTGAAGAGCTCTCTGTTCTCTGCGACGCCGATGTTGCTCTCATCATCTTCTCTTCCACCGGAAAAC
SVP_O_msk_CDS   TCTTTTCAAGAAAGCTGAAGAACTCTCTGTTCTCTGCGACGCCGATGTTGCTCTCATGATCTTCTCTTCCACCGGAAAGC

...............................................................................................

                161                                                                           240
SVP_R_msk_CDS   TCTTCGA---------------CATGAAGGAAGTATTAGAGAGGCATAACTTGCAGTCAAAGAACTTGGAGAAGCTTGAT
SVP_O_msk_CDS   TCTTCGAGTTCTGTAACTCCAGCATGAAGGAAGTATTAGAGAGGCATAACTTGCAGTCAAAGAACTTGGAGAAGCTTGAT

...............................................................................................

                241                                                                           320
SVP_R_msk_CDS   CAGCCATCTCTTGAGTTACAGCTGGTTGAGAACAGTGATCACGCCCGAATGAGTAAAGAAATTGCCGACAAGAGCCACAG
SVP_O_msk_CDS   CAGCCATCTCTTGAGTTACAGCTGGTTGAGAACAGTGATCACGCCCGAATGAGTAAAGAAATTGCCGACAAGAGCCACAG

...............................................................................................

                321                                                                           400
SVP_R_msk_CDS   ACTAAGGCAAATGAGAGGAGAGGAGCTTCATGGACTTAACATTGAAGAGCTGCAACAGCTAGAGAAAGCCCTCGAAGCTG
SVP_O_msk_CDS   ACTAAGGCAAATGAGAGGAGAGGAGCTTCAGGGACTTAACATTGAAGAGCTGCAACAGCTAGAGAAAGCCCTCGAAGCTG

...............................................................................................

                401                                                                           480
SVP_R_msk_CDS   GTTTGACCCGCGTGATTGAAACAAAGAGTGACAAGATTATGAATGAGATTAGCGAACTTCAAAGAAAGGGAATGCAATTG
SVP_O_msk_CDS   GTTTGACCCGCGTGATTGAAACAAAGAGTGACAAGATTATGAATGAGATTAGCGAACTTCAAAGAAAGGGAATGCAATTG

...............................................................................................

                481                                                                           560
SVP_R_msk_CDS   ATGGATGAGAACAAGCGGTTGAGGCAGCAAGGAACTCAACTAACGGAAGAGAACGAGCGACTTGGCATGCAAATATATAA
SVP_O_msk_CDS   ATGGATGAGAACAAGCGGTTGAGGCAGCAAGGAACTCAACTAACGGAAGAGAACGAGCGACTTGGCATGCAAATATATAA

...............................................................................................

                561                                                                           640
SVP_R_msk_CDS   TAACGTGCATGAATCCGGTGTTGATGAATCGGAGAACGTCGCCGTGTACGAGGAAGGACACTCGTCAGAGTCTATTACTA
SVP_O_msk_CDS   TAACGTGCATGAATCCGGTGTTGATGAATCGGAGAACGTCGCCGTGTACGAGGAAGGACACTCGTCAGAGTCTATTACTA

...............................................................................................

                641                                                                           720
SVP_R_msk_CDS   ACGCCGGAAACTCCAACGGAGCTCCTGTTGACTCCGAGAGCTCCGACACTTCCCTTAGGCTCGGCTTACCGTATGGTGGT
SVP_O_msk_CDS   ACGCCGGAAACTCAAACGGAGCTCCTGTTGACTCCGAGAGCTCCGATACTTCCCTTAGACTCGGGTAA------------

...............................................................................................

                721
SVP_R_msk_CDS   TAG
SVP_O_msk_CDS   ---

...............................................................................................
```

**Fig.S4** Locations of the selected spacers in the first exons of *CO* (a) and *SVP* (b) homoeologs. For each homoeolog sgRNAs used in the study are indicated with blue color below the amino acid sequences, adjacent PAM sites are marked with orange. Below gRNA annotations 1st exons of CO and SVP genes are marked with yellow, and whole gene marked with light green, red highlighted nucleotides mark mismatches between homoeolog sequences. Figures were created from screenshots of MAFFT alignments made in the Benchling web-service (https://www.benchling.com/)

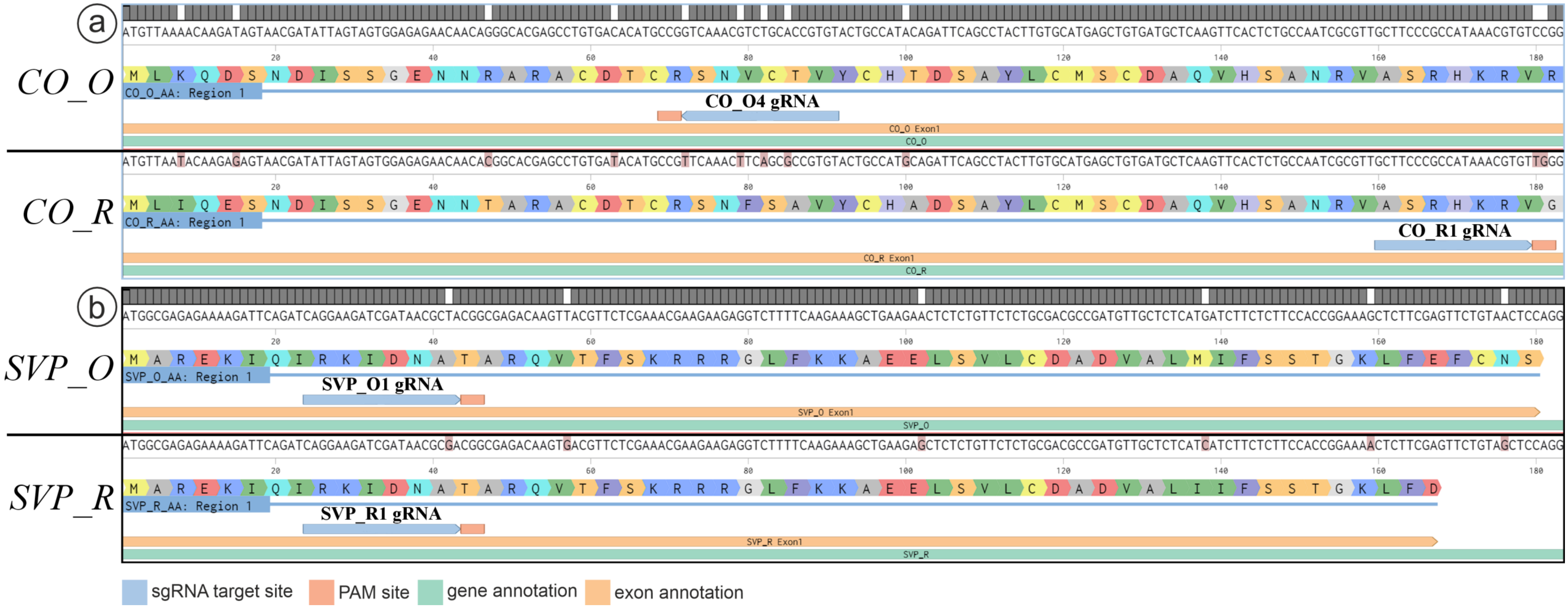
